# Epigenetic aging in brain tissue of the self-fertilizing vertebrate, *Kryptolebias marmoratus*

**DOI:** 10.1101/2025.09.19.677260

**Authors:** Justine Bélik, Frédéric Silvestre

## Abstract

DNA methylation changes predictably with age across taxa, but in most species these patterns are confounded by genetic variation. As a result, age-predictive methylation models have mostly been developed in genetically heterogeneous, cross-fertilizing organisms, limiting inference about epigenetic aging per se. Disentangling epigenetic and genetic effects is therefore essential for understanding aging, adaptation, and evolution. Here, we exploit the mangrove rivulus (*Kryptolebias marmoratus*), one of only two known self-fertilizing vertebrates (together with *K. hermaphroditus*), to examine epigenetic aging in a system of naturally occurring near-isogenic individuals. Using reduced-representation bisulfite sequencing of 90 brain samples spanning 60-1100 days of age, we identified 40 CpG sites whose methylation levels predict chronological age with high accuracy (R^2^ > 0.96, Median Absolute Error of 28.7 days). These 40 age-associated CpG sites were linked to nearby genes with known roles in cellular maintenance and neurodegeneration. These include genes implicated in aging and neurodegenerative processes across vertebrates, such as lamin-A, the aryl hydrocarbon receptor, and genes associated with Alzheimer’s disease in humans. By leveraging a self-fertilizing vertebrate, this study demonstrates that DNA methylation undergoes consistent, age-associated changes across the lifespan in the near absence of genetic variation. Our results establish self-fertilizing vertebrates as powerful models for disentangling epigenetic aging from genetic effects and provide a foundation for comparative and evolutionary studies of aging.

**STATEMENTS:** *Data availability:* The datasets generated and/or analyzed during the current study are available in the NCBI repository, under the ID *BioProject ID PRJNA1331489*.

*Funding statement:* This work was supported by the FNRS project J.0189.24 “Epigenome Stability in Mangrove Rivulus”.

*Ethics approval:* All research reported in this manuscript was conducted in accordance with institutional and national ethical standards for animal care and use. Experimental procedures involving *Kryptolebias marmoratus* were approved by the Animal Experimentation Ethics Committee (UN PM KE 23/020).

*Authors’ contributions:* JB designed the experimental process, conceptualized the article, generated and analyzed the data and wrote the manuscript. FS designed the experimental process, reviewed and edited the manuscript and did validation and supervision. All authors read and approved of the final manuscript.

*Conflict of interest:* The authors declare that they have no competing interests.

*Patient consent statement:* Not applicable.

*Permission to reproduce material from other sources:* Not applicable.

*Clinical trial registration:* Not applicable.

## 1. INTRODUCTION

Aging is a progressive, time-dependent decline in physiological integrity, leading to impaired function and an increased risk of death among most living organisms (Jones et al., 2015; López-Otín et al., 2013; Rose et al., 2012). It is considered the primary risk factor for several major human pathologies, including cancer, diabetes, cardiovascular disorders and neurodegenerative diseases, highlighting its central role in ongoing scientific research. López-Otín et al. (2013) initially described nine “Hallmarks of Aging”: genomic instability, telomere attrition, epigenetic alterations, loss of proteostasis, deregulated nutrient sensing, mitochondrial dysfunction, cellular senescence, stem cell exhaustion, and altered intercellular communication. These hallmarks have been widely studied and validated. However, the list has since been updated to include disabled macroautophagy, chronic inflammation, and dysbiosis (López-Otín et al., 2023). Each hallmark represents a distinct yet interconnected entry point for investigating the biology of aging, as they collectively influence and reinforce one another.

Among the hallmarks of aging, epigenetic alterations play a significant role. These include changes in DNA methylation patterns, abnormal posttranslational modifications of histones, aberrant chromatin remodeling, and deregulated functions of noncoding RNAs (López-Otín et al., 2023). Such modifications can profoundly impact gene expression, contributing to the onset and progression of several age-related pathologies. In particular, the age-associated accumulation of DNA methylation (DNAm) modifications has been extensively studied (Seale et al., 2022). DNAm involves the addition of a methyl group to a cytosine, most commonly at CpG dinucleotides in vertebrates (i.e., a cytosine located 5′ of a guanine). This process is achieved by a set of enzymes from the DNA methyltransferase (DNMT) family, notably, DNMT1 (which copies preexisting DNA methylation motifs onto the new strand during mitosis) and DNMT3a/3b (which perform *de novo* methylation). DNAm can be reversed through two primary processes: active demethylation, which is mediated by ten-eleven translocases (TETs) dioxygenases, and passive demethylation, which occurs when functional DNMT activity is absent during DNA replication (Davletgildeeva & Kuznetsov, 2024).

Current literature demonstrates that aging is strongly correlated with DNAm, with substantial functional consequences. Early studies suggested a global hypomethylation with aging (Fuke et al., 2004; Singhal et al., 1987; Wilson et al., 1987) but more recent studies produced conflicting results and do not support this hypothesis (Fasolino et al., 2017; Lister et al., 2013; Raddatz et al., 2013). Either way, DNAm is both gained and lost with aging, resulting in a more variable DNAm pattern in older individuals (Meyer & Schumacher, 2024). This includes hypermethylation of previously unmethylated regions and hypomethylation of previously methylated regions, increasing the overall entropy of the DNAm pattern over time (Meyer & Schumacher, 2024; Tarkhov et al., 2024; Tong et al., 2024). However, some CpG sites exhibit a strong, consistent correlation with chronological age, emphasizing their potential as age predictors (De Paoli-Iseppi et al., 2017). These alterations are often tissue- and species-specific (Unnikrishnan et al., 2018). These so-called “clock CpGs” form the basis of epigenetic clocks, sets of carefully selected loci across the genome that are collectively capable of predicting chronological age with high accuracy (Piferrer & Anastasiadi, 2023).

Key epigenetic clocks include the model developed by Horvath (Horvath, 2013), constructed from 8,000 human multi-tissue samples, which uses 353 CpGs to estimate age with a median error of 3.6 years. The Hannum clock (Hannum et al., 2013), which is based solely on human blood samples, highlights specific components of human aging. Levine et al. (Levine et al., 2018) advanced the concept of biological age *versus* chronological age by incorporating clinical health biomarkers into the clock’s design. In parallel, numerous epigenetic clocks have been built across the tree of life. In mammals, examples include *Mus musculus* (Spiers et al., 2016; Stubbs et al., 2017) and *Canis lupus familiaris* (Thompson et al., 2017). In birds, clocks have been constructed for *Gallus gallus* (Gryzinska et al., 2013) and *Coturnix japonica* (Andraszek et al., 2014). Clocks also exist for reptiles, such as *Alligator mississippiens* (Nilsen et al., 2016), and fish, including *Danio rerio* (Mayne et al., 2020), the European seabass (Piferrer et al., 2024) and the haddock (Strand et al., 2025). In addition, the first epigenetic clocks for invertebrate species are now emerging (Brink et al., n.d.; Guynes et al., 2024). The recent development of a universal pan-mammalian clock suggests that epigenetic aging is evolutionarily conserved across mammals (Lu et al., 2023). Similarly, a study on cetaceans introduced a multi-species, two-tissue clock, highlighting the potential to develop tools applicable across diverse taxa (Zoller et al., 2025).

The further development of epigenetic clocks is valuable, as it offers insights into aging mechanisms and related diseases. Comparative studies can identify universal and species-specific aspects of aging and epigenetic regulation (Bertucci-Richter & Parrott, 2023). Moreover, several independent studies have demonstrated that epigenetic age is indicative of past and/or present health status (Caulton et al., 2022). Epigenetic clocks offer yet underexploited potential in the context of breeding programs and fisheries management (Piferrer & Anastasiadi, 2023), as well as conservation work (Heydenrych et al., 2021). Age estimation notably enables the calculation of growth rates (to assess species productivity and growth variation over time), mortality rates (natural and fishing mortality rates) and population age structure (Piferrer & Anastasiadi, 2023).

Most epigenetic clocks are based on easily accessible tissues, such as blood or caudal fins, to enable nonlethal and minimally invasive age estimation (Piferrer et al., 2024; Piferrer & Anastasiadi, 2023). However, these tissues may not be the most relevant for increasing our fundamental understanding of aging. In contrast, investigating the aging brain methylome is critically important, given its influence on neural function and the development of neurodegenerative diseases (Liu et al., 2009; Pellegrini et al., 2021; Thrush et al., 2022; Yang et al., 2015). Only a few epigenetic clocks have been constructed via postmortem brain samples, either in nonhuman primates (Horvath et al., 2021; Jasinska et al., 2021) or humans (Grodstein et al., 2021; Shastri et al., 2024). To our knowledge, only two studies have focused on the brain in the context of the epigenetic clock (Coninx et al., 2020; Stubbs et al., 2017). Thus, expanding our knowledge of brain-specific methylation patterns in the context of aging has become an essential objective for both epigenetic and neuroscience research.

Here, we focus on the mangrove rivulus, *Kryptolebias marmoratus*. With its sister species, *Kryptolebias hermaphoditus*, they are the only known self-fertilizing vertebrates (Costa et al., 2010). This unique feature allows to study naturally clonal lineage and disentangle epigenetic and genetic effects. *K. marmoratus* belongs to a broad group of killifish species and is distributed along the coasts of Florida to northern Brazil, closely following the red mangrove *Rhizophora mangle* (Taylor, 2012). Its reproduction strategy is remarkable, as it involves a rare mixed mating system known as androdioecy (Costa et al., 2010). Populations of *K. marmoratus* are composed of self-fertilizing hermaphrodites, which produce clonal offspring after a few generations, and males, which can cross-fertilize unfertilized eggs laid by hermaphrodites. In natural populations, there is a gradient in male frequency, resulting in varying levels of genetic diversity, with selfing rates ranging from 49 to 97% (Chapelle, 2023). The population used in our study originates from Emerson Point Preserve, Florida, and is characterized by an extremely high selfing rate of 97% and an observed heterozygosity of zero. The virtual absence of genetic variation offers a unique opportunity to study epigenetic variation without the confounding effects of underlying genetic differences, and is essential to understand aging, adaptation and evolution.

We analyzed 90 brain samples from hermaphroditic *Kryptolebias marmoratus* individuals of known age and spanning the species’ lifespan (60 days to 3.5 years post-hatching). Using reduced-representation bisulfite sequencing (RRBS), we identified 40 CpG sites whose combined methylation profiles accurately predict chronological age (Median Absolute Error, 28.7 days; R^2^ = 0.96). Genomic annotations of these sites revealed associations with nearby genes involved in aging-related processes, including DNA damage repair, mTOR signaling, inflammation, and autophagy, as well as genes implicated in Alzheimer’s disease in human studies. Together, these results demonstrate that age-associated DNA methylation changes arise consistently across the lifespan of a near-isogenic vertebrate, providing evidence that epigenetic aging dynamics can be detected largely independently of genetic variation.

## 2. MATERIAL AND METHODS

### 2.1 Mangrove rivulus aging colony

*Kryptolebias marmoratus* individuals were collected in 2019 from Emerson Point Preserve, Florida (N25°01’45.64”, W80°29’49.24”, permit number SAL-24-1132**A**-SR) and used to establish the F0 generation at the University of Namur. The fish were individually housed in 500 mL plastic containers filled with 300 mL of 12 ± 1 parts per thousand (ppt) saltwater (Instant Ocean™ Sea salt) and maintained in a climate-controlled room at 25 ± 1°C under a 12:12 light:dark photoperiod. The fish were fed daily *ad libitum* with live *Artemia salina*.

For the epigenetic clock analysis, we selected 96 hermaphrodite individuals, covering the major life span of the mangrove rivulus (from 60 to 1100 days). The age of the individuals is known to the day, as they hatched in the laboratory. The number of fish was decided following Mayne et al. (2021), knowing that our species is self-fertilizing, which reduces genetic and therefore epigenetic variability, and that we use only one sex. The fish were between the 2^nd^ and 6^th^ generations derived from the F0 stock (field individuals). Out of the 96 fish selected, four were excluded after being identified as secondary males, and two were removed because of insufficient DNA yield. Once they reached the expected age, the fish were euthanized by rapid cooling (immersion in water between 0°C and 4°C for 20 seconds), and death was confirmed by decapitation. Each individual was photographed, measured and weighed. Detailed information on individual fish is provided in Supplementary Table S1. All procedures were approved by the animal ethics committee of the University of Namur (UN PM KE 23/020). Following euthanasia, brain, liver, gonad and fin tissues were collected and stored at -80°C until DNA extraction.

### 2.2 Reduced-Representation Bisulfite Sequencing of brain samples

#### 2.2.1 DNA extraction

DNA was extracted using NucleoSpin Tissue XS kit (Marcherey-Nagel, item number 740901.250), with slight modifications to the manufacturer’s protocol. An additional wash step, identical to the first, was included, and a 5-minute incubation at 37°C was added following the binding buffer step. Elution was performed using 2x10 µL (instead of once 20 µL), with each followed by centrifugation at 11,000 × *g* for 1 minute. DNA quality and quantity were assessed via both a NanoDrop spectrophotometer and a high-intensity Qubit fluorimeter. Samples that did not meet the minimum amount of DNA required for the Diagenode RRBS kit (< 50 ng of DNA, see below) were excluded from further analyses (two samples in total).

#### 2.2.2 Reduced-Representation Bisulfite Sequencing

RRBS libraries were prepared via the Premium RRBS Kit V2 from Diagenode (catalog number C02030037) following the manufacturer’s protocol. This kit includes all the reagents required for enzymatic digestion, library preparation, bisulfite conversion and PCR amplification. The final DNA concentrations were determined via a high-sensitivity Qubit fluorimeter. Libraries were sequenced on a NovaSeq 6000 V1.5 (300 cycles XP workflow) with paired-end sequencing (20 million reads each way) via the GIGA platform (University of Liège, Belgium). The FASTQ files were quality checked with FastQC v0.11.8 (https://www.bioinformatics.babraham.ac.uk/projects/fastqc/).

### 2.3 Bioinformatics

#### 2.3.1 Methylation analysis

RAW FASTQ files were trimmed using *Trim Galore!* (version 0.6.10) with the following parameters: -rrbs - quality 28 -illumina -stringency 2 -length 40. Genome indexing and bisulfite alignment were performed using *Bismark* (v. 0.24.1). The latest version of the reference genome of *K. marmoratus* was used (GCF_001649575.2_ASM164957v2) and prepared via *Bismark_genome_preparation*. Read alignment was conducted via *Bowtie2* through *Bismark* with the following parameters: -score_min L,0, 0.6. *SAMtools* (version 1.17) was then used to sort and index the resulting BAM files.

The files were then processed in R (version 4.2.2) via the *methylKit* (version 1.28.0) and *caret* (version 6.0-94) packages. Bam files were processed with *processBismarkAln* via assembly = ASM164957v2 and read.context = “CpG”. Coverage filtering was applied via *filterByCoverage*, which retained only sites with a minimum coverage of 15 and maximum coverage below 99.9^th^ percentile to mitigate PCR bias.

#### 2.3.2 Predicting age from CpG methylation

The model was built and validated following the main steps shown in **Figure 1**. The CpGs present in all the samples were first selected using *methylKit::unite. percMethylation* function was used to create the percentage methylation matrix. The first selection of CpGs was carried out by correlating each CpG with the logarithm of the age of the individuals (*cor*.*test*, with Benjamini-Hochberg correction). Age was transformed to a natural log to fit a linear model. Preselecting the CpG stabilizes the sites selected in the final model when several seeds are compared. Preselection of informative features in high-throughput datasets has been shown to improve model performance (Newediuk et al., 2025). By eliminating uninformative predictors at an early stage, feature selection reduces overfitting and enhances prediction accuracy on independent datasets (Theng et al., 2025).

**Figure 1.**
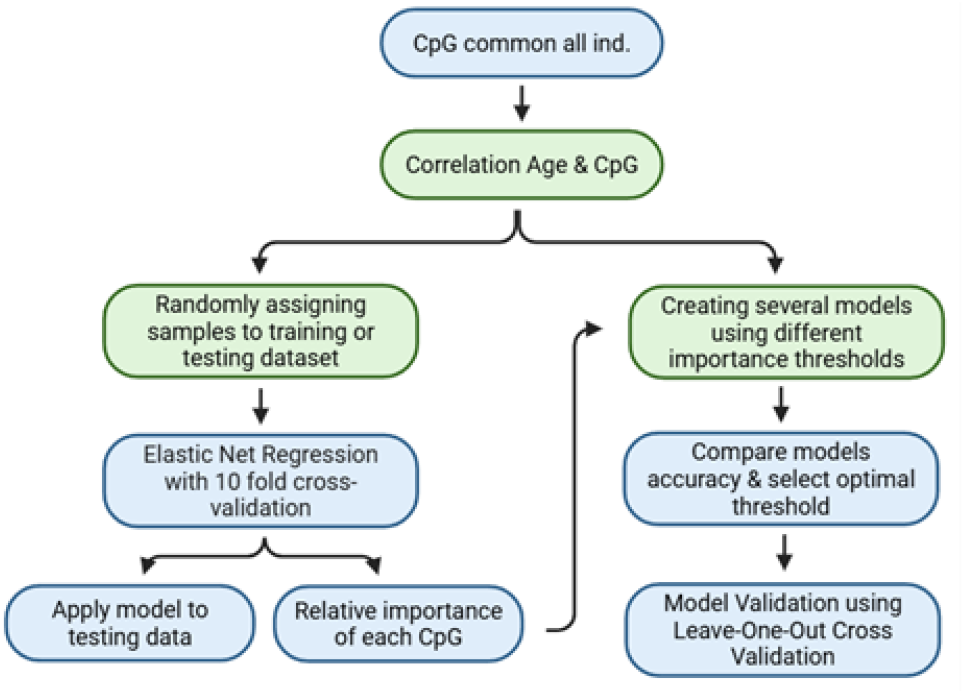
diagram of the main stages in building and validating the model. The green frames are steps that have been studied and optimized. Their conclusions are discussed in the materials and methods section.

The samples were randomly assigned to either a training (80%) or a testing dataset (20%) via the *createDataPartition* function in the caret R package. This division is necessary to prevent false improvements in model predictions. We tested the most common division, 70-30, 75-25 and 80-20, and compared the model accuracy with three different seeds. The 80–20 partition always provided the best results (Table S2).

The *training* function from the *caret* package was used with 10-fold cross-validation to perform elastic net regression on the training dataset. The use of elastic net regression is optimized for high throughput data as the hyperparameters α and λ shrink uninformative sites to zero and remove them for the models (Kuhn & Johnson, 2013). *caret* tests the combination of three α (0.1, 0.5 and 1) and λ (depending on the data, always between 0 and 1) values and selects the combination best suited to the data, i.e., the one with the highest R^2^ (strength of linear relationship between chronological and epigenetic age) and lowest Median Absolute Error (MAE, precision of the clock’s predictions) on the testing data.

To focus on biologically informative markers of brain aging, we further reduced the number of CpG sites by selecting an optimal trade-off between model complexity and predictive performance. The elastic net model assigns each CpG a relative importance score (0–100), which we used to iteratively filter predictors. We first conducted a coarse screening using an importance threshold ranging from 5 to 75 in steps of 5, followed by a finer screening between 25 and 35 in steps of 1. The final model was selected based on achieving high predictive accuracy (R^2^ > 0.95) with the smallest number of CpGs, when evaluated on the testing dataset. This stepwise approach revealed a non-linear relationship between the number of predictors and model performance, indicating that subsets of CpGs contribute unequally and are not strictly complementary.

The initial model was trained using all 90 individuals, yielding an R^2^ of 0.937 and a mean absolute error (MAE) of 35.6 days (Fig. 2A). Because prediction accuracy decreased in older individuals, we repeated the entire modelling procedure using only the 76 individuals younger than 900 days. This age-restricted model showed improved performance (R^2^ = 0.96; MAE = 28.7 days; Fig. 2B) and was therefore retained as the final epigenetic clock. Unless stated otherwise, all subsequent results and interpretations are based on this reduced model.

**Figure 2.**
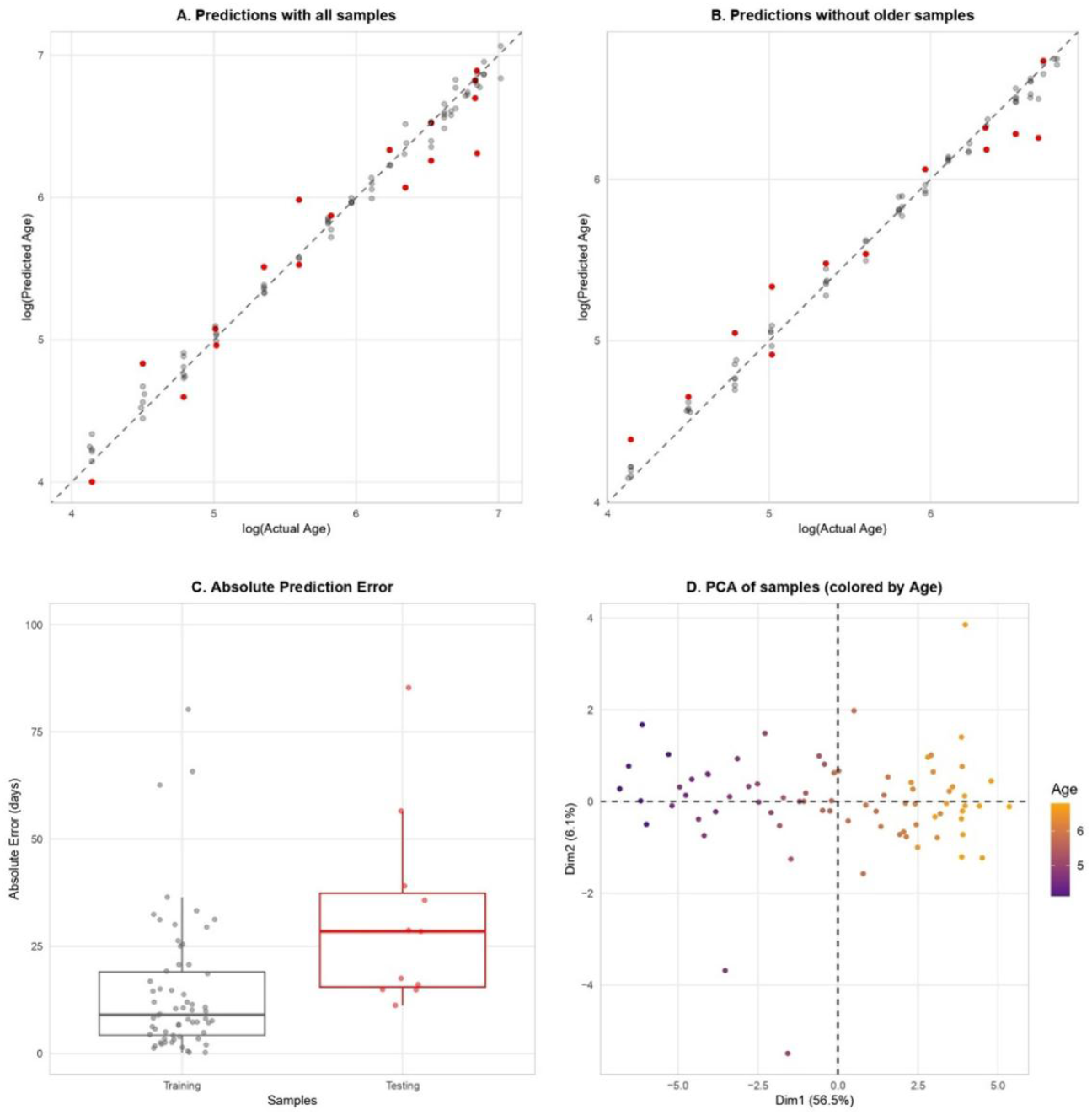
Mangrove rivulus age estimation. Chronological age is plotted against predicted age, for both training (in gray) and testing (in red) datasets, for both all individuals (A) and individuals younger than 900 days only (B). Absolute Error for both training and testing datasets, for the regression excluding the older individuals, with Welch-test (C). PCA of individuals using the 40 CpG sites, showing a clear separation based on age in the first principal component (D).

The model was ultimately evaluated using Leave-One-Out Cross Validation (LOOCV), where each fold consists of a single observation. In this approach, the model is trained on all data points except one, which is then used for testing. This process is repeated systematically so that every observation serves once as the test set. LOOCV provides an estimate of the true test error, as training is always performed on the full dataset minus one sample (James et al., 2013). The results revealed no difference between the model’s performance and its validation.

PCA was then performed on methylation data using the *FactoMineR* package (version 2.10), to visualize the individuals with respect to their age.

The code is available here: https://github.com/jubelik/Epigenetic-clock.

### 2.4 Annotation

A file containing the chromosome, start and stop positions, and strand information for each selected CpG site was imported into *SeqMonk* (version 1.481) and analyzed using the genome assembly ASM164957v2 (Kelley et al., 2016). Probes were generated by assigning CpG sites to genes located within a maximum distance of 20,000 base pairs. The resulting annotation file included, in addition to the imported data, the associated gene name (*Features*), the Feature Orientation (upstream, overlapping, downstream, or null), and the distance between the CpG site and the gene when not overlapping. The roles of the genes were described via Z-fins to maintain consistency in the annotations.

## 3. RESULTS

### 3.1 Age-associated CpG sites identified by RRBS

Information about the fish used (Fish ID, Age at death and Length) are in Table S1. The bisulfite conversion was 98.87%. RRBS samples were aligned to the *K. marmoratus* reference genome, with an average alignment rate of 56.8%. The percentage of global methylation was 60.2%. Global methylation did not significantly correlate with age (p value = 0.5748, Figure S1).

A total of 467,182 CpG sites were common to all individuals, with a minimum coverage of 15 and maximum coverage set at the 99.9th percentile. Among these sites, 36,704 CpGs were significantly correlated with age (correlation test, corrected for multiple testing via the Benjamini-Hochberg procedure). The selected regression, using 80/20 split, has a α value of 0.1 and a λ value of 0.05. This model achieved an R^2^ of 0.938 and MAE of 35.6 days, using 38 CpG. However, as the correlation was less accurate in older individuals (**Figure 2A**), a second model was rerun using only individuals younger than 900 days (76 individuals). A total of 475,502 CpG sites were common to these individuals, with a minimum coverageof 15 and maximum coverage set at the 99.9th percentile. Among these sites, 29,126 were significantly correlated with age (correlation test, corrected for multiple testing via the Benjamini-Hochberg procedure). The selected regression has a α value of 0.1 and a λ value of 0.150. The final model included 40 CpG sites, achieved an R^2^ of 0.960 and MAE of 28.7 days (**Figure 2B**).

Validation of the model by Leave-One-Out-Cross-Validation (LOOCV) yielded virtually identical results, indicating that both approaches capture true accuracy equally well. A MAE of 28.7 days was found in the testing dataset, and no statistical difference was observed between the absolute error rate between the training and testing data sets (p-value = 0.066, welch-test, two-tailed) (**Figure 2C**).

We subsequently performed a PCA of the individuals based on their methylation profile (**Figure 2D**), with chronological age represented as a color gradient. The first principal component explained 56.5% of the total variance and was strongly associated with age. The second component explained 6.1% of the total variance.

### 3.2 Identification and characterization of age-related CpG sites

Age-associated DNA methylation levels are often highly correlated across CpG sites. In the presence of multicollinearity, elastic net regression select one predictor arbitrarily from a set of correlated CpGs, leading to variability in the specific sites retained across repeated model fits (Engebretsen & Bohlin, 2019). Accordingly, annotation of individual CpG sites included in the final model should not be interpreted as uniquely informative or causal (Moqri et al., 2023), but rather to provide genomic context.

Information for each CpG site and its associated gene is summarized in **Table 1**. Among the 40 selected CpG sites, 31 showed an increase in the percentage of methylation with age, whereas 9 exhibited a decrease. Among these sites, 31 were located within gene bodies, 8 were positioned upstream (ranging from 232 to 5,622 base pairs from the start codon), and 1 was located downstream (11,307 base pairs). Each gene was evaluated for methylation level changes (DNAm change), strength of association (R^2^), genomic context, distance to the gene if not within it, and functional relevance (Table 1). The age-associated methylation trajectories of individual CpG sites display substantial heterogeneity and are shown in Figure S2.

**Table 1:**
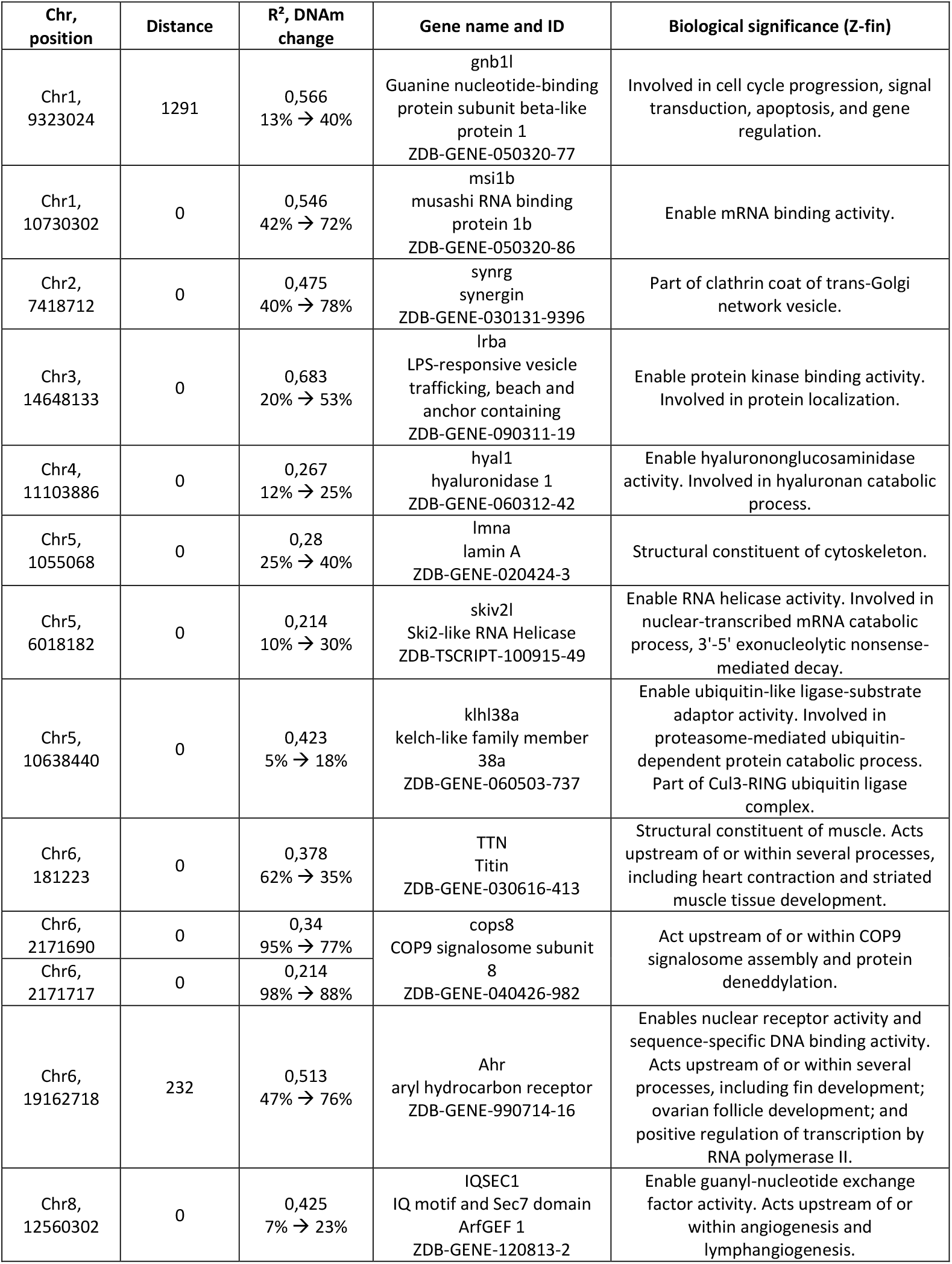
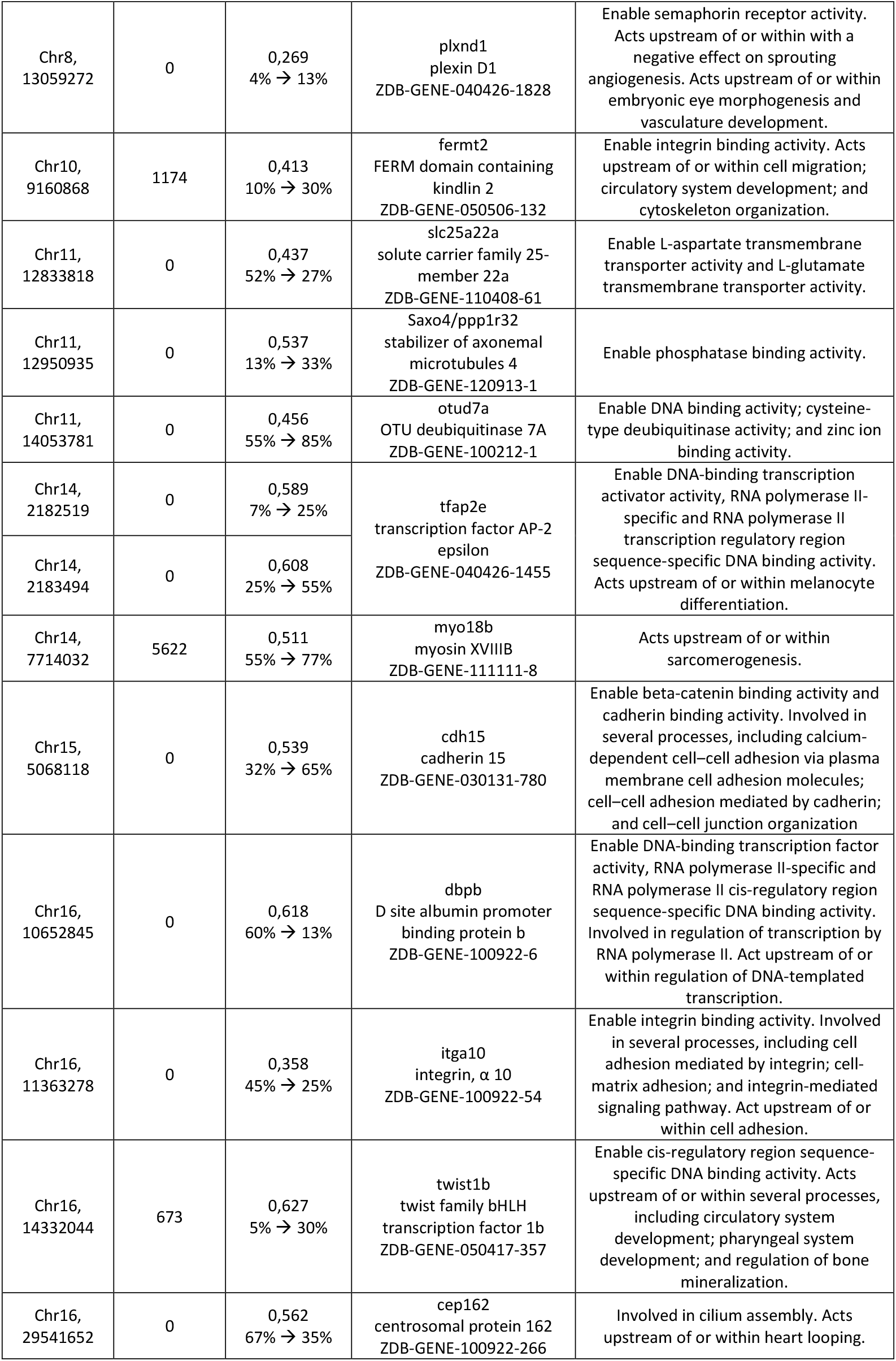
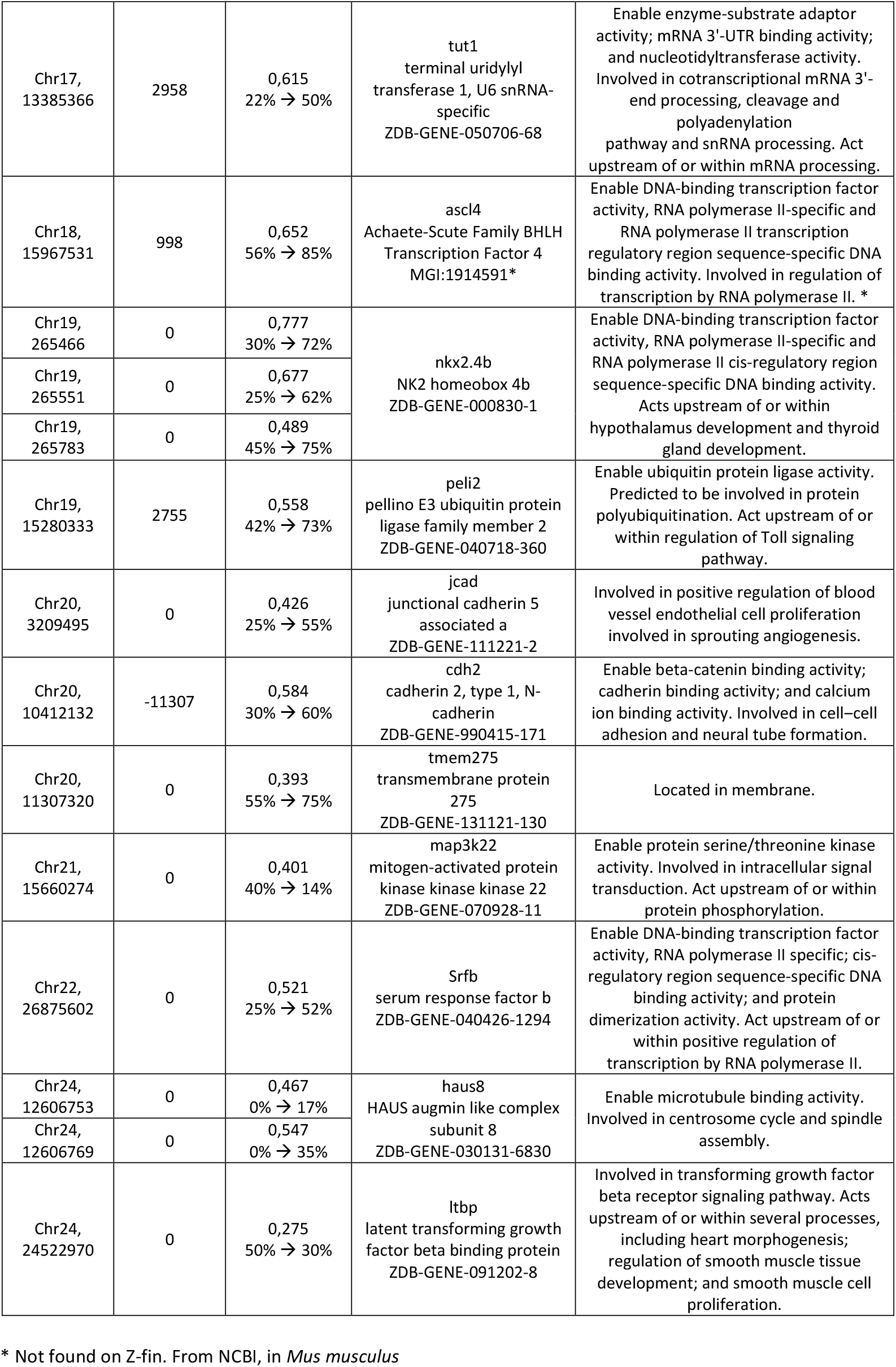
Chromosome and position, Distance to the gene, R^2^ and DNA methylation change (from 60 DPH to 900 DPH), Gene name and ID and Biological Significance using Z-fin.

The CpG sites with the highest R^2^ values are within the genes nkx2.4b (position 265466, R^2^ of 0.777), lrba (R^2^ of 0.683), nkx2.4b (position 265551, R^2^ of 0.677), ascl4 (R^2^ of 0.652) and dbpb (0.618). Their methylation levels changed by at least 30%, with an increase for the first four and a decrease for dbpb (60% to 13%). The nkx2.4b gene appears twice in the 5 most correlated CpG, and a third CpG site present in this gene was also selected within our 40 CpG sites, with an R^2^ of 0.489. Two CpG sites were also selected within the same gene for cops8 (R^2^ values of 0.340 and 0.214, both decreasing methylation), haus8 (R^2^ values of 0.547 and 0.467, both increasing methylation), and tfap2e (R^2^ values of 0.608 and 0.589, both increasing methylation).

Among the genes found, several essential functions were identified. Notably, cell cycle progression, signal progression and apoptosis (gnb1l), mRNA binding activity (msi1b), mRNA 3’-end processing, cleavage and polyadenylation (tut1), protein kinase activity (lrba), cytoskeleton (lmna), structural component of muscle (titin), microtubule binding activity (haus8), DNA binding transcription activity (tfap2e, dbpb, nkx2.4b and srfb), serine/threonine kinase activity (map3k22) and transformation of the growth factor beta receptor signaling pathway (ltbp) were investigated. The complete list of gene roles associated with the selected CpG sites can be found in Table 1.

## 4. DISCUSSION

In this study, we developed the first brain-based epigenetic clock for a teleost fish. Using the mangrove rivulus *Kryptolebias marmoratus*, we investigated non-pathological brain aging in a system characterized by extremely low genetic variation (Taylor, 2000). *K. marmoratus* exhibits an androdioecious mating system composed of self-fertilizing hermaphrodites and males, generating naturally occurring near-isogenic lineages. Self-fertilization dramatically reduces genetic variation among individuals, providing a powerful framework to disentangle age-associated epigenetic changes from genetic background effects (Kelley et al., 2016). These features make *K. marmoratus* a uniquely suited model for studying epigenetic aging with minimal genetic confounding.

Beyond constructing an epigenetic age predictor in a self-fertilizing vertebrate, we aimed to characterize baseline epigenetic aging dynamics in the brain. We identified 40 CpG sites whose combined methylation profiles predict chronological age with high accuracy (R^2^ > 0.96; mean absolute error = 28.7 days). Among genes associated with the selected CpGs, several have attracted our attention in the study of aging. It is important to underline that the selected CpGs should not be interpreted as uniquely determining aging-related processes or as having direct functional roles (Moquri et al., 2023). Gene annotations associated with these CpG sites are therefore presented to provide biological context and to facilitate comparison with aging-related pathways described in other systems, rather than to imply a causal relationship between methylation at individual loci and gene function. Nevertheless, these associations provide a foundation for future experimental studies to test causal relationships between specific methylation changes and aging phenotypes.

The CpG at position 1055068 on chromosome 5 is within the gene *lamin-A* (*lmna*) and shows hypermethylation with age (from 22% of methylation at 60 days to 40% at 900 days). *lmna* encodes the prelamin-A/C protein, which undergoes four post-translational steps to become mature lamin A, the main constituent of the nuclear lamina (Lin & Worman, 1993). Defect in the post-translational maturation disrupt nuclear function and affect aging-related processes, such as mTOR signaling, epigenetic modifications, the stress response, inflammation, microRNA activation and mechanosignaling, causing premature aging syndromes such as Hutchinson–Gilford progeria syndrome (Cenni et al., 2020). Moreover, lamin A, which is typically not expressed in neurons, appears to transform from senile to AD neurons and contributes to halting the consequences of cell cycle re-entry and nuclear Tau exit, allowing the survival of the neuron. This irreversible nuclear transformation leads to neurofibrillary tangle formation and, ultimately, neurodegeneration (Gil et al., 2021).

A second CpG site of interest is located at position 19162718 on chromosome 6, 232 base pairs upstream from the gene *Aryl Hydrocarbon Receptor* (AhR). It shows hypermethylation with age, ranging from 47% to 77%. AhR is a ligand-dependent transcription factor that is classically associated with the regulation of xenobiotic metabolism in response to environmental toxicants (Hankinson, 1995). Several AhR ligands, such as resveratrol and quercetin, have been shown to extend the lifespan of model organisms (Bhullar & Hubbard, 2015; Kampkötter et al., 2008; Pietsch et al., 2009). However, conflicting results have been obtained, depending on the ligand, species, and experimental setup used (Eckers et al., 2016; Serna et al., 2024). It has also been suggested that AhR reveals evidence of antagonistic pleiotropy in the regulation of the aging process, i.e., the gene is beneficial during development but shows deleterious properties during aging (Salminen, 2022).

Two selected CpG sites are within *cops8* (COP9 signalosome 8) and are both hypomethylated with age (position 2171690, ranging from 98% methylation to 77%, and position 2171717, ranging from 98% to 87%). The COP9 signalosome participates in several cellular and developmental processes in various eukaryotic organisms and is associated with de-ubiquitination and protein kinase activities (Wei & Deng, 2003). Subunit 8 is a central regulator of the proteasomal proteolytic pathway and selective autophagy (Su et al., 2011). The autophagic–lysosomal pathway is critical for the removal of oxidized proteins in the heart (Su et al., 2013). Like epigenetic alterations, impaired autophagy is one of the primary hallmarks of aging (López-Otín et al., 2023).

Among the other selected CpGs, several show a link with AD. A Genome Wide Association study revealed that FERM domain containing kindlin 2 (*fermt2)* (CpG 1174 base pair upstream, which is upregulated with age) is a locus susceptible to AD. These results suggested that *ferm2* modulates AD risk by regulating APP metabolism and Aβ peptide production (Chapuis et al., 2017). Another GWAS revealed that an increased risk of developing AD has been associated with 27 genes, including fermt2. LPS-responsive vesicle trafficking, beach and anchor containing (*Lrba*), solute carrier family 25-member 22a (*slc25A22*) and integrin α 10 (*itga10*) are differentially expressed in AD patients compared with healthy cognitive adults and have been found in our epigenetic clock (Kim et al., 2023; Tian et al., 2024; Zou et al., 2023).

Our study identified molecular markers of baseline brain aging in the mangrove rivulus that are also implicated in age-related neurological diseases in humans. This overlap highlights the potential of experimentally perturbing these markers to assess their causal roles in pathological aging. Such perturbations could be achieved through controlled exposure to neurotoxic compounds or via targeted genetic and epigenetic manipulations, including CRISPR–Cas9 and CRISPR–dCas9–based approaches (Cai et al., 2023; Caobi et al., 2020; Chen et al., 2018; Ruetz et al., 2024). Although we have focused on CpGs associated with genes that have been linked in literature to aging and/or neurodegeneration, it is important to study the mechanism of other CpGs in the future. A particular focus should be placed on the nkx2.4b, haus8 and tfap2e genes, for which several CpGs have been selected. These genes may have an underestimated role in the aging and death of individuals, but no literature has mentioned them. Exploring these candidates not only broadens our understanding of potential contributors to aging but also provides an opportunity to investigate how their alteration may drive or protect against disease processes.

The selected CpG sites and the sequencing data generated add to the data already available for other teleost species, e.g., European sea bass (Anastasiadi & Piferrer, 2020), scorpion fish (Weber et al., 2022), cow nose rays (Weber et al., 2024) and haddock (Strand et al., 2025), all of which show very high accuracy. The increasing number of studies on the teleost epigenetic clock has increased interest in the construction of a pan-teleost epigenetic clock, which would be particularly valuable in the fields of aquaculture and conservation (Piferrer & Anastasiadi, 2023). A first pan-mammalian epigenetic clock have been built, including 185 species and 59 tissue types, and show very high accuracy (r > 0.96) (Lu et al., 2023). This pan-epigenetic clock proposes solutions to solve the inconstancy in sequencing methods, different lifespans, etc., underlying the feasibility of such a teleost epigenetic clock. Other prospects include applying this epigenetic clock to other mangrove rivulus lineages, with various genetic diversity, as well as natural populations. This method would provide valuable ecological information by enabling the estimation of age structures in natural populations, thereby deepening our understanding of demographic processes and life history traits (Piferrer & Anastasiadi, 2023).

Many epigenetic clocks have been described in the literature; however, there remains little consensus regarding key regression parameters used in their construction. In this study, we compared different modeling choices to assess their influence on predictive performance within our dataset. Notably, the 80-20 split between training and testing data consistently yielded superior performance on the testing dataset (Table S2). To avoid the risk of overfitting (artificially inflating model performance), we encouraged implementing a 10-fold cross-validation procedure, which provides a more robust estimate of model generalizability by evaluating performance across multiple training–testing splits within the training data. Furthermore, we encourage a first selection of the predictor based on their own correlation with age, as it reduces model complexity and improves stability (Engebretsen & Bohlin, 2019). Prioritizing CpG with stronger individual age association improve interpretability and better reflect selected biological processes involving specific epigenetic features (Bertucci & Parrott, 2020). Regarding the final CpG selection, we noticed a non-linear relationship between the number of CpG sites included and predictive accuracy, therefore we recommend a careful evaluation of the inclusion threshold to achieve a parsimonious model with stable performance.

As has been seen in other work (e.g. El Khoury et al., 2019), our epigenetic clock tends to underestimate chronological age in older individuals. One hypothesis for this bias is early saturation of methylation levels at selected CpG sites, such that the regression model extrapolates to biologically implausible methylation values. Another contributing factor may be the presence of 5-hydroxymethylcytosine (5hmC), which cannot be distinguished from 5-methylcytosine (5mC) using conventional bisulfite sequencing and may distort age predictions in tissues with high 5hmC abundance, such as the brain (El Khoury et al., 2019). Although the level of 5hmC in mangrove rivulus brain tissue has not yet been characterized, studies in other teleost fish indicate that hydroxymethylation plays an important regulatory role. In *Oreochromis niloticus*, differential hydroxymethylation has been reported in genes associated with growth, particularly within gene bodies and promoters (Konstantinidis et al., 2023). Age-associated epigenetic drift may further contribute to reduced predictive accuracy in older individuals (Hannum et al., 2013; Heyn et al., 2012).

In conclusion, our study not only revealed age-related changes in CpG methylation patterns but also represents the first development of an epigenetic clock in a self-fertilizing vertebrate, the mangrove rivulus. This unique model allows the study of DNAm changes without the confounding effects of genetic variation, providing insights into the specific roles of DNAm in aging. Using brain tissue enabled us to investigate baseline aging processes in this organ and to gain functional insights into brain aging. The genes identified here warrant further investigation to improve our understanding of brain aging and its links to neurodegenerative diseases.

## ACKNOWLEDGEMENTS

We thank our lab technician Jérôme Lambert for their help with the library preparation. We also thank Alice Dennis for proofreading the article.

This work was supported by the FNRS project J.0189.24 “Epigenome Stability in Mangrove Rivulus”.

## Notes

### Competing Interest Statement

The authors have declared no competing interest.

### Summary of Updates

The paper has been improved in terms of its writing. The major changes concern the discussion and its narrative flow. The methods and results have not changed.

