## Supplementary materials for "Epigenetic aging in brain tissue of the self-fertilizing vertebrate, *Kryptolebias marmoratus*"

**Table S1: Identity (ID), age at death (days), and length (mm) of the fish used.**

| **Fish ID** | **Age (days)** | **Length (mm)** | **Fish ID** | **Age (days)** | **Length (mm)** | **Fish ID** | **Age (days)** | **Length (mm)** |
| --- | --- | --- | --- | --- | --- | --- | --- | --- |
| EPP23-57 | 62 | 17,72 | EPP23-35 | 212 | 29,97 | EPP21-183 | 683 | 37,47 |
| EPP23-52 | 63 | 19,42 | EPP23-27 | 270 | 29,2 | EPP21-184 | 683 | 35,91 |
| EPP23-53 | 63 | 19,76 | EPP23-28 | 270 | 32,24 | EPP21-118 | 748 | 39,07 |
| EPP23-54 | 63 | 18,85 | EPP23-29 | 270 | 32,1 | EPP21-142 | 749 | 37,25 |
| EPP23-55 | 63 | 18,86 | EPP23-30 | 270 | 31,11 | EPP21-143 | 749 | 36,4 |
| EPP23-56 | 63 | 18,93 | EPP22-49 | 331 | 36,44 | EPP21-146 | 749 | 38,71 |
| EPP23-50 | 89 | 21,23 | EPP22-50 | 331 | 37,2 | EPP21-147 | 749 | 38,64 |
| EPP23-46 | 90 | 20,48 | EPP22-51 | 331 | 35,64 | EPP21-27 | 786 | 43,45 |
| EPP23-47 | 90 | 22,06 | EPP22-43 | 338 | 34 | EPP21-28 | 786 | 41,65 |
| EPP23-48 | 90 | 20,89 | EPP22-44 | 338 | 36,55 | EPP21-111 | 811 | 37,55 |
| EPP23-49 | 90 | 21,37 | EPP22-20 | 388 | 38,98 | EPP21-112 | 811 | 36,8 |
| EPP23-45 | 91 | 22,37 | EPP22-37 | 390 | 37,57 | EPP21-113 | 811 | 38,54 |
| EPP23-59 | 120 | 25,19 | EPP22-38 | 390 | 35,95 | EPP21-105 | 869 | 34,98 |
| EPP23-60 | 120 | 25,46 | EPP22-39 | 390 | 37,93 | EPP21-68 | 882 | 39,61 |
| EPP23-61 | 120 | 23,85 | EPP22-45 | 390 | 37,48 | EPP21-71 | 882 | 44,09 |
| EPP23-62 | 120 | 26,46 | EPP22-18 | 449 | 37,07 | EPP21-108 | 924 | 40,87 |
| EPP23-58 | 121 | 24,89 | EPP22-11 | 450 | 36,34 | EPP21-63 | 930 | 45,55 |
| EPP23-63 | 121 | 25,44 | EPP22-12 | 450 | 37,02 | EPP21-64 | 930 | 42,0 |
| EPP23-64 | 121 | 31,6 | EPP22-14 | 450 | 36,73 | EPP21-101 | 931 | 35,57 |
| EPP23-41 | 150 | 26,08 | EPP22-03 | 510 | 35,06 | EPP21-99 | 931 | 38,98 |
| EPP23-42 | 150 | 25,83 | EPP22-05 | 510 | 37,72 | EPP21-102 | 942 | 38,4 |
| EPP23-39 | 151 | 26,16 | EPP22-01 | 513 | 37,26 | EPP21-154 | 944 | 39,57 |
| EPP23-40 | 151 | 27,24 | EPP22-13 | 566 | 38,4 | EPP21-168 | 944 | 39,47 |
| EPP23-43 | 151 | 26,7 | EPP22-10 | 570 | 39,05 | EPP21-169 | 944 | 39,82 |
| EPP23-44 | 151 | 24,41 | EPP22-28 | 570 | 40,54 | EPP21-116 | 961 | 43,2 |
| EPP23-32 | 211 | 28,9 | EPP22-19 | 574 | 40,81 | EPP21-98 | 989 | 39,65 |
| EPP23-33 | 211 | 28,81 | EPP21-176 | 683 | 37,73 | EPP21-39 | 990 | 44,83 |
| EPP23-36 | 211 | 25,46 | EPP21-178 | 683 | 40,28 | EPP21-40 | 990 | 44,6 |
| EPP23-37 | 211 | 30,35 | EPP21-180 | 683 | 37,74 | EPP20-18 | 1114 | 46,79 |
| EPP23-38 | 211 | 30,54 | EPP21-182 | 683 | 38,19 | EPP20-20 | 1114 | 48,26 |

**Table S2: Comparison of the 70–30, 75–25, and 80–20 divisions on 3 different seed, with the number of CpG and respective R². When no cut-off offers exactly 40 CpGs, the number of CpGs was chosen to be as close to 40 as possible.**

| Seed | Division | #CpG | R² |
| --- | --- | --- | --- |
| 935 | 70-30 | 40 | 0,852 |
|  | 75-25 | 39 | 0,938 |
|  | 80-20 | 40 | **0,960** |
| 123 | 70-30 | 41 | 0,920 |
|  | 75-25 | 38 | 0,893 |
|  | 80-20 | 42 | **0,964** |
| 2 | 70-30 | 41 | 0,859 |
|  | 75-25 | 41 | 0,916 |
|  | 80-20 | 40 | **0,955** |

**Figure S1: Global methylation level with age.**

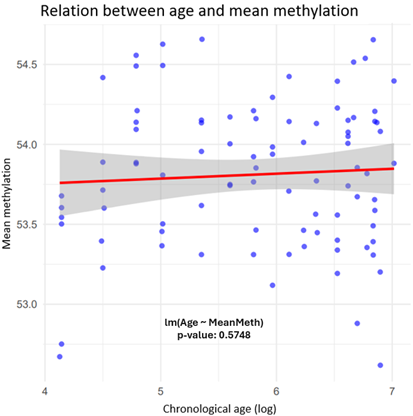

**Figure S2: DNA methylation level of 40 selected CpG sites across chronological age.**

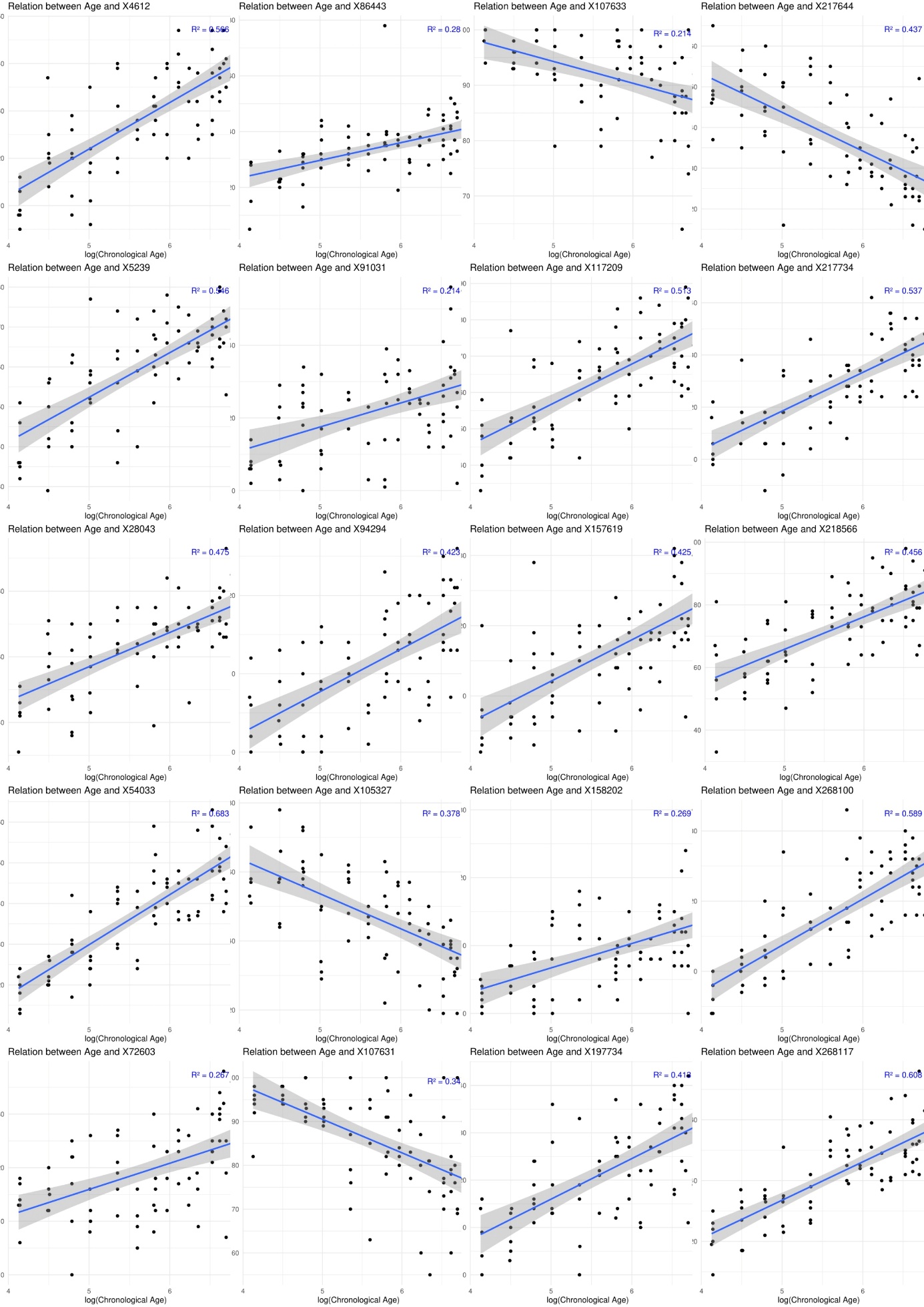

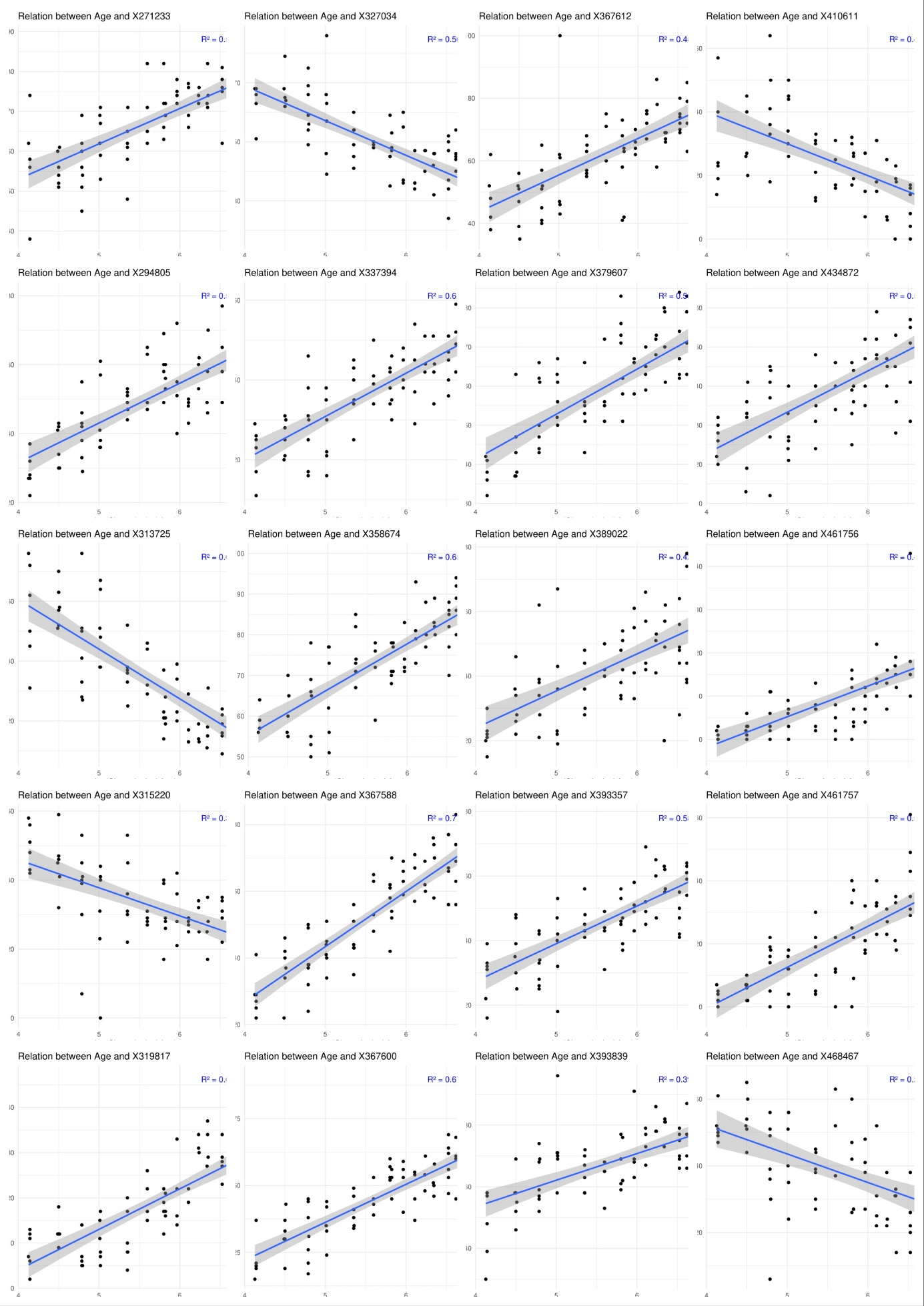
